# A new species of Lepanthopsis (Orchidaceae, Pleurothallidinae) in the Atlantic Forest in São Paulo, Brazil

**DOI:** 10.1101/703694

**Authors:** L.R. Zandoná, E.L.M. Catharino, A.G. Maragni, M.M.F. Jesus

## Abstract

A new species of Lepanthopsis is discovered and described for the Atlantic Forest of southeastern Brazil, the species called *Lepanthopsis legadensis* is distinguished from other Brazilian species, notably by the petals and long-acuminate sepals.

## Introduction

Orchidaceae surveys in the Atlantic Forest of the state of São Paulo reveal a high species richness, despite the current state of degradation of these forests, notably studies carried out in Conservation Units in recent years, such as the State Park of Cardoso Island (147 species, Romanini And Barros 2008), in Mogi das Cruzes, Chiquinho Veríssimo Municipal Park (70 species, Rodrigues & Barros 2012) and still higher than that found in Serra do Japi in Jundiaí (125 species, Pansarin & Pansarin 2008) of Ipiranga – PEFI (125 species, Barros 1983). This number was also higher than that found in Juréia-Itatins Ecological Station in the Juréia Massif (77 species, Catharino & Barros 2004) and Cantareira State Park (159 species, Zandoná & Catharino 2015). The latter revealed the largest number of species occurring in a delimited area of the Atlantic Forest, many endemic or threatened with extinction, such as *Centroglossa macroceras*, *Grandiphyllum hians*, *Grandiphyllum divaricatum*, which until then were considered as extinct in nature. This last study was based on a survey and non-predatory collections, with periodic rescue in fallen trees, cultivation and later herborization based on the plants rescued and kept in cultivation.

An ongoing study, which is part of the research project: Orchids of Legado das Águas, conservation of rare and endangered species, started in November 2015, reveals, after 238 days of field visits, the presence of 221 species of orchids, 13 in the red list of endangered flora of the State of São Paulo (São Paulo 2016), using the same non-predatory methodology adopted by Zandoná & Catharino 2015. (Zandoná et.al. 2019 unpublished data).

Many of the collected species are still under study but already demonstrating the presence of rare and threatened species. In the course of the identification of the rescued and flowering species in cultivation, we come across an interesting species of Lepanthopsis (Figs. 1 and 2) Cogn. (Ames). This genus, which is little represented in Brazilian flora, now has more than 40 recognized species distributed in tropical forests of the Americas (Luer 1991, Pridgeon 2005). Only three of these species are registered in Brazil. Two of them, *L. densiflora* and *L. floripecten* have been known for a long time and occur in Atlantic forests of southern and southeastern Brazil (Hoehne 1949, Pabst and Dungs 1975, 1977), and a third, *L.vellozicola*, occurs as endemic in a region of rupestrian fields in Minas Gerais (Mota et.al. 2009).

**Figure 1.**
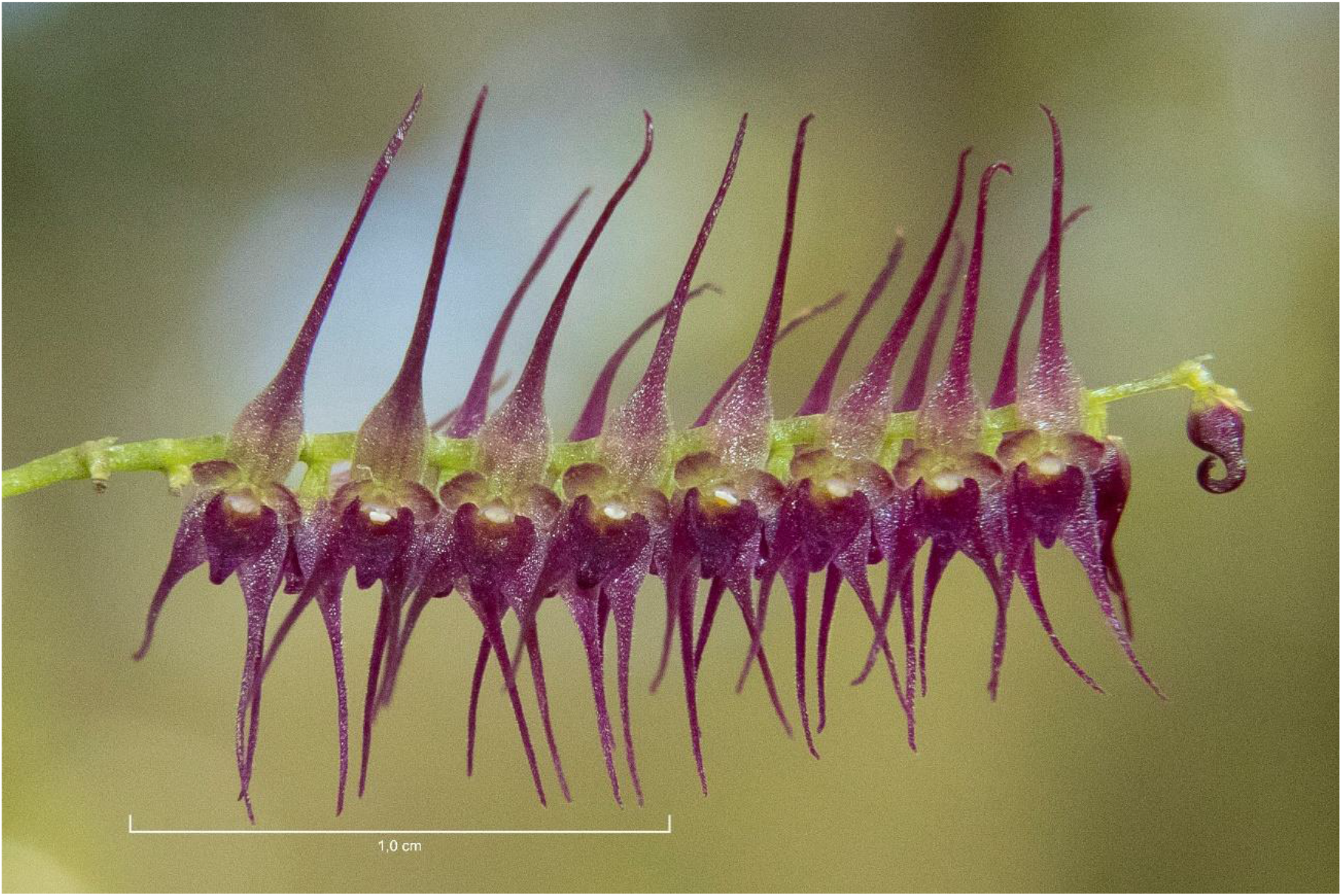
*Lepanthopsis legadensis* Zand.& Cath. Photo Luciano Zandoná (15.04.2018)

**Figure 2.**
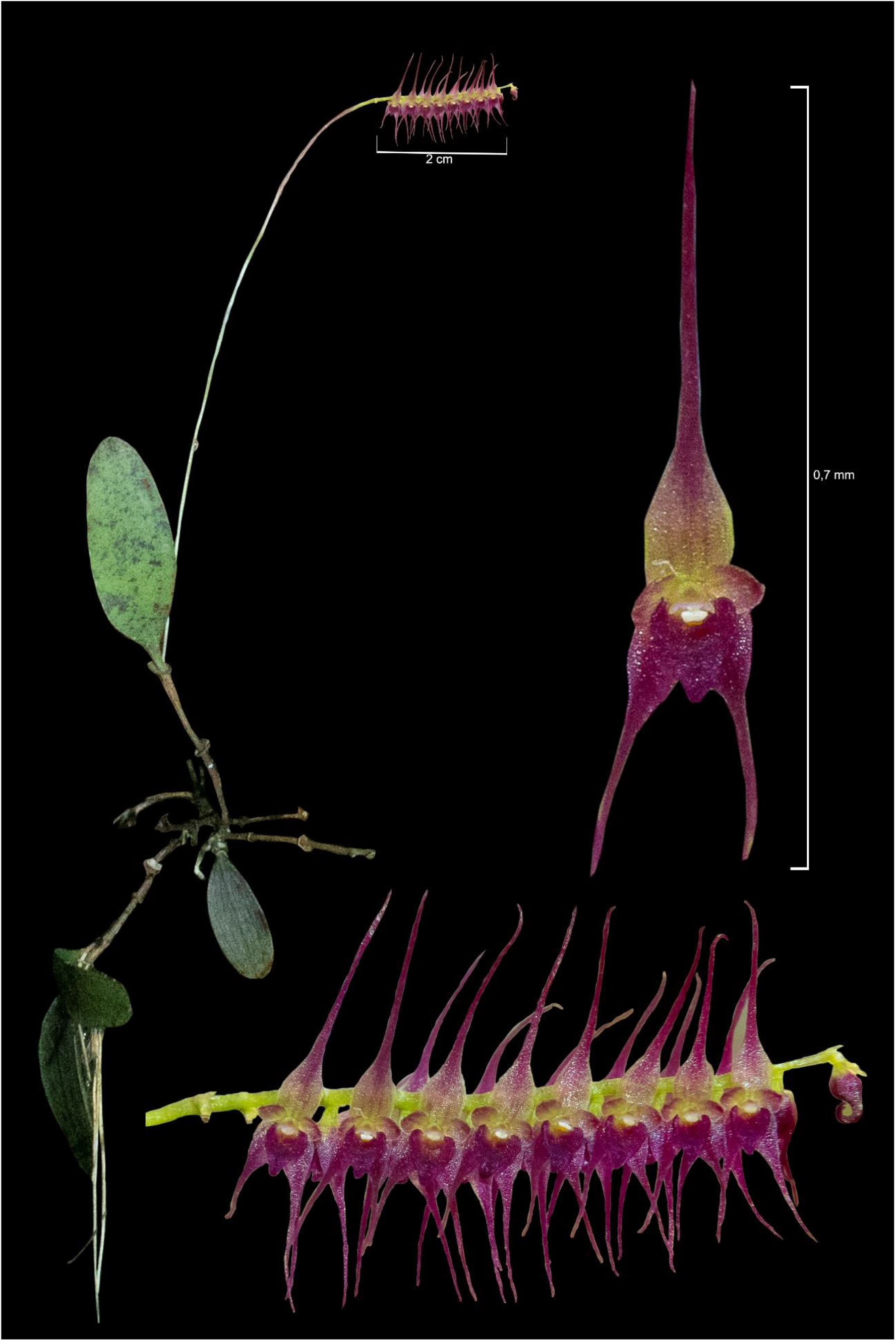
*Lepanthopsis legadensis* Zand. & Cath.,Photographic plate mounted from photos of the plant in cultivation. Photos: Luciano Zandoná

By consulting the relevant bibliographies and materials deposited in the Herbarium SP and also the SpeciesLink network, in an attempt to identify this *Lepanthopsis* specimen, it was concluded that this is a new and remarkable species for science, included in the subgenus Lepanthopsis (Luer 1991), which is described in this paper, with an updated key presentation of Lepanthopsis species for Brazil.

## Taxonomy

### Lepanthopsis legadensis

L.R. Zandoná & E.L.M. Catharino, ***sp.nov***. (Figs.1 e 2)

Type: BRAZIL. São Paulo: Miracatu, Legado das Águas, coord: 262415 E 7340519 S, 18/07/2017 (Bloom in green house in 15/04/2018), Zandoná, L.R., Zandoná, A.G.Maragni & Jesus, M.M. F. de (holotype, SP 498030).

Small epiphyte plant; Roots ca. 6cm, not very numerous, relatively thick; Ramicauls ca. 4 cm, wrapped in 3 lepantiform sheaths with smooth margins; Leaves erect, coriaceous ca. 3.0×0.9 cm, oblong to obovate, rounded apex and cuneate base, attenuated in pseudo-petiole with ca. 3 to 4 mm, irregular irregular stains on the abaxial face; Inflorescence in racemous congestion, with long peduncle, total length of ca. 8.5 cm, with about 20 flowers distributed in the 2 cm apical of the peduncle, opening almost simultaneously in two opposite rows; Floral bracts ca. 0.2mm; Flowers vinaceous, membranous sepals, long-acuminate, with evident central rib, Dorsal sepal with ca. 5 mm long, 1 mm wide at base, yellowish in this region; Lateral sepals ca. 4 mm long, 1 mm wide at the base, practically free from each other, lightly conated at the base only; Petals membranous, semi-orbicular, concave, ca. 0.7×07mm, glandular-ciliated; Lip slightly chordate, ca. 1,1mm length and ca. 1mm wide, papillary and slightly glandular-ciliated, apex sub-acute to obtuse, deflexus, obtuse lateral lobes embracing the spine, disc of the centrally concave lip and with yellowish spot in the central region, Column ca.0,4mm of compr. and broad, broader at apex; Anther apical, Bilobed apical stigma. Pollinia 2.

## Diagnostic characteristics

The color, the size of the stem, the long acuminate sepals, the lateral ones being practically free from each other and contained only at the base, are marked characteristics of the species described herein. The sepals and long-acuminate petals differentiate this species from all other Brazilian ones. Although it does not occur in Brazil, *Lepanthopsis acuminata* Ames is apparently the closest to the species described, which can however be differentiated by the size of the smaller plant, the longest flower stem in relation to the plant, the sepals cones in the base in a larger proportion that the present species, the cordiform lip, being in the species described here.

## Etymology

Dedicated to the Legado das Águas, Atlantic Forest reserve located in the Vale do Ribeira, belonging to the Votorantim Group, where the plant was collected.

## Distribution, habitat and phenology

The species described is only known in a small area of Legado das Águas, located in the Ribeira de Iguape Valley, between the municipalities of Miracatu and Tapirai, State of São Paulo, Brazil. The “Legado” is the largest private reserve in the Atlantic Forest, with 31,000 ha of high conservation forests, mostly primitive forests. The species was located in the high of 650 to 700m, vegetating in the inner canopy to about 15m of height, along with rare *Octomeria estrellensis*.

The Lepanthopsis described above was rescued in a fallen tree in July 2017, without flowers, only with two dried flower stems, and included in the living collection of Legado das Águas as *Lepanthopsis* sp. and kept in cultivation. It flowered in April 2018, when it was discovered that it was a new species. A ramicauls was herborized according to the conventional techniques described in Fidalgo & Bononi (2010), and also some flowers preserved in alcohol 70% and deposited in Herbarium SP (SP 498030).

## Simplified and updated key of Brazilian Lepanthopsis species

1. 1. lateral sepals, only found at the base, long-acuminate sepals…. ***L. legadensis*** Lateral sepals cones at least half the length, sepals not arched….**2**
2. Fully conical side sepals…..***L. velloziana*** Partially conical side sepals, from the base to at least half of its length…**3**
3. Side sepals cones half or close to it, apex of acute lateral sepals, ovate labellum… ***L. densiflora*** Lateral sepals conated up to near apex, apex of obtuse lateral sepals, suborbicular lip…***L. floripecten***

## IUCN Red List Category

According to current knowledge and based on IUCN Red List Categories and Criteria (IUCN, 2012), Lepanthopsis legadensis can be considered critically endangered (CR), with only two individuals located in an area less than 1km2 in mature forest.

## Acknowledgements

The authors would like to thank all the team of Legado das Águas and especially Ms. Frineia Rezende da Silva and mr. David Canassa responsible for Votorantim Reserves. They also thank the Votorantim Group.

## Notes

#### Summary of Updates

references updated; Species description updated

